# Quantifying Exposure of Pacific Salmon and Steelhead to Climate Change in the Fraser River basin

**DOI:** 10.64898/2026.03.18.712684

**Authors:** Stephanie J. Peacock, William Cheung, Brendan Connors, Lisa Crozier, Sue Grant, Eric Hertz, Brian P.V. Hunt, Josephine C. Iacarella, Cory R. Lagasse, R. Dan Moore, Jonathan W. Moore, Francois Nicolas-Robinne, Marc Porter, Markus Schnorbus, Samantha M. Wilson, Katrina Connors

## Abstract

Climate change can affect salmon and steelhead (*Oncorhynchus spp.*) throughout their anadromous life cycles, yet there have been no assessments of which Canadian populations face the greatest exposure. We developed a framework to quantify relative climate change exposure of salmon and steelhead populations based on the spatial and temporal distribution of different life stages. Exposure was calculated from climate model projections for freshwater and marine climate variables considering unique impact thresholds for each population and life stage. We applied this framework to 60 Conservation Units of Pacific salmon and steelhead in the Fraser River basin, British Columbia. Lake-type sockeye had the highest exposure, driven by elevated stream temperatures during adult freshwater migration and spawning stages and relatively low thermal tolerance of marine stages. Chinook salmon were the next most exposed, while coho, pink, and chum salmon had relatively low exposure. Uniquely, steelhead exposure was driven by high stream temperatures during incubation. Our framework is broadly applicable, and our findings provide critical input for climate change vulnerability assessments and forward-looking resilience planning for Pacific salmon.

## Introduction

Anadromous fishes are particularly susceptible to the impacts of climate change because they integrate these effects across their freshwater and marine life-stages (McDaniels et al. 2010; Waldman and Quinn 2022; Crozier and Siegel 2023). Many Pacific salmon (*Oncorhynchus spp.*) and steelhead (*O. mykiss*) (henceforth, “salmon”) populations have declined throughout their western North American range in response to decades of cumulative anthropogenic pressures, including climate change (Munsch et al. 2022; Connors et al. 2024). As recovery and rebuilding efforts expand, there is a pressing need to understand and account for the challenges and opportunities that climate change presents for salmon - now and into the future (Moore and Schindler 2022; MacDonald and Grant 2023). However, quantitative assessments of future conditions that salmon will be exposed to throughout their lifecycle are limited – especially in Canada (but see McDaniels et al. 2010; Iacarella and Weller 2023). Without information on climate change exposure, the effectiveness of recovery efforts may be undermined and strategies to reduce risks to biodiversity and maintain salmon resilience will remain elusive.

One challenge to predicting how climate change will affect salmon is that their exposure, and responses, to climate changes are mediated by diverse life-history traits. Salmon tend to return to their natal rivers to spawn, allowing for local adaptations to their freshwater habitats to evolve over time. These adaptations include life-history traits such as migration and spawn timing and the duration of juvenile freshwater rearing (Beechie et al. 2006). Ocean entry timing and early marine distributions also differ among salmon populations and are a response to local marine conditions (e.g., Wilson et al. 2023). Diversity in life histories, genetics, and habitat use among salmon in Canada is represented by Conservation Units (CUs): groups of salmon that are sufficiently genetically and ecologically unique that they are unlikely to be recovered within a human lifetime if extirpated (Fisheries and Oceans Canada 2005). The differences in traits among CUs have been well-documented in some cases - for example, Fraser sockeye CUs are assigned to different river-entry timing groups that are closely monitored to inform in-season fisheries management (e.g., Michielsens and Cave 2019). Only recently, however, has life-history timing for CUs been compiled into a centralized repository (Wilson and Peacock 2025) that can be used to quantify how this diversity influences the relative exposure of salmon to climate change.

Another common limitation to understanding exposure for aquatic species like salmon is the availability of climate change projections for relevant environmental variables at the appropriate spatial and temporal scales corresponding to each life-stage. In the marine environment, statistical downscaling of coarse global models is needed to understand differences in exposure arising from differential use of estuarine habitats by CUs. In freshwater, stream temperature projections are limited, and although proxies like air temperature and precipitation are more readily available, they may not relate well to river conditions experienced by aquatic species. For example, rising air temperatures may result in increasing or decreasing stream temperature depending on whether river input is dominated by rainfall or glacial melt. Different approaches have been developed to estimate stream temperature and flow from other global climate variables. Statistical models of August mean temperatures fitted to historical data (e.g., Isaak et al. 2017; Weller et al. 2023) have recently allowed for a spatial comparison of projected warming and habitat suitability for salmon at a relatively fine spatial resolution (Iacarella and Weller 2023). However, these models lack the temporal resolution to examine how different life-history timings may impact the relative exposure of CUs. Hydrologic models that incorporate mechanistic relationships among drivers are more flexible to project variables throughout the year and are well suited to making long-term projections of how salmon freshwater habitats will change. Downscaling of projections from multiple Global Climate Models (GCMs) for British Columbia (BC), Canada using mechanistic hydrologic models has recently been expanded to include output for stream temperature and flow at a daily time step (Schnorbus 2024). This advance allows a more temporally fine-scaled consideration of how life-history timing impacts exposure. Further, these hydrologic projections are available for the same individual GCMs as marine climate projections, providing a more consistent view of model uncertainty in overall exposure across the life-cycle (Wade et al. 2017; Crozier et al. 2021).

Here, we develop and apply a framework for quantifying the climate change exposure of salmon. We define climate change exposure as the proportion of time that a CU is expected to experience adverse environmental conditions in future periods, considering both the magnitude of physical climate changes projected to occur and the sensitivity of CUs to those changes based on attributes such as geographic distribution, migration timing, and thermal tolerance. Our framework for assessing exposure is quantitative and data-driven, making it scalable and transferable to other regions or species. We apply the framework to assess climate change exposure for 60 Pacific salmon and steelhead CUs in the Fraser River basin, British Columbia. This information is a critical first step to help support forward-looking and proactive strategies to increase the resilience of salmon in the face of climate change.

## Methods

### Assessment units

We focused our assessment of climate change exposure on Pacific salmon and steelhead CUs (Fisheries and Oceans Canada 2005), of which there are 60 in the Fraser River basin. Conservation Units reflect differences in habitat use, genetics, and ecology among groups of salmon, and are thus well-suited to understanding risks to biodiversity, but they are generally finer than scales of fisheries management (e.g., Pacific Fisheries Management Areas or Stock Management Units). Conservation Units also capture the within-species variations in life-history traits that will affect exposure to climate changes. For example, Chinook CUs tend to have different freshwater residence time, ocean residence time, or adult freshwater migration timing.

Conservation Units have been defined by Fisheries and Oceans Canada (DFO) for Chinook (*O. tshawytscha*), chum (*O. keta*), coho (*O. kisutch*), pink (*O. gorbuscha*), and sockeye (*O. nerka*) salmon (Holtby and Ciruna 2007). We include six steelhead CUs that have been proposed by the Province of BC (Parkinson et al. 2005; Salmon Watersheds Program 2024), and are based on similar principles. A complete list of the 60 CUs and their life-history characteristics is included in the Online Supplement (Table S1).

### Life stages

We took a life-cycle approach to assessing exposure that considers climate changes in both freshwater and marine environments. For each CU, we defined six life stages based on life-history timing, spatial distribution, and sensitivity to environmental conditions:

1. Incubation, when eggs are developing in the gravel;
2. Freshwater rearing, when juvenile salmon have hatched and may move within connected freshwater habitats;
3. Early marine, when salmon enter the ocean and spend a critical period of growth and development in near-shore waters;
4. Marine rearing, when salmon transition from the coast to offshore for a period of one to six years;
5. Adult freshwater migration, when salmon re-enter rivers on their return migration to the spawning grounds; and
6. Spawning, when salmon are mating and laying eggs on the spawning grounds.

We considered four major life-history transitions that dictated the timing (start and duration) of each stage. The **spawning** period marked both the extent of the spawning stage and the start of the incubation stage. The transition from incubation to juvenile rearing occurred with **fry migration**, when young salmon emerge from the gravel after their development through the egg and alevin stages. **Ocean entry**, when juvenile salmon migrate from freshwater to the marine environment, marked the transition between juvenile rearing and early marine stages. **Freshwater re-entry** (i.e., run timing), when adult salmon return to rivers enroute to spawning grounds, marked the end of the marine rearing stage and start of the adult freshwater migration. Based on these life-history transitions, we defined the timing and duration of the six life stages such that there was some overlap between stages in order to capture the variability in life-history timing within a CU (Table 1).

**Table 1.** Exposure to climate changes was calculated for each of six life stages based on timing and spatial distribution.

| Stage | Timing | Spatial distribution |
| --- | --- | --- |
| Incubation | Start* of **spawning** to end of **fry migration** | Accessible stream reaches with optimal gradient for spawning for the given species (Norris 2024) within the CU boundary |
| Freshwater rearing | Start of **fry migration** to end of **ocean entry** | Accessible stream reaches with optimal gradient for rearing for the given species (Norris 2024) within the CU boundary, or known rearing lake for lake-type sockeye CUs |
| Early marine | 3 months before to 3 months after peak **ocean entry** date | A 300 to 750 km buffer zone around point of ocean entry |
| Marine rearing | 3 months after **ocean entry** to end of **freshwater re-entry** | Pacific Ocean (north of 35°N) and Bering Sea |
| Adult freshwater migration | Start of **freshwater re-entry** to start of **spawning**<sup>†</sup> | Migration route(s) from river entry point to accessible spawning stream reaches |
| Spawning | Start to end of **spawning** period | Accessible stream reaches with optimal gradient for spawning for the given species (Norris 2024) within the CU boundary |

\* Start, peak, and end dates correspond to the average 2.5<sup>th</sup>, 50<sup>th</sup>, and 97.5<sup>th</sup> quantiles of timing distributions for the CU; see Wilson and Peacock (2025) for details.
†To avoid unrealistically long exposure times for early migrating lake-type sockeye, the adult freshwater migration stage for these CUs was shortened to the expected duration of migration to the holding lake; see Online Supplement.

The timings of these transitions were compiled for each of the 60 CUs by Wilson and Peacock (2025). For CUs without observations, we estimated timing from available data. Data gaps were most prevalent for the fry migration and ocean entry transitions, which we imputed using the weighted mean of timings from CUs of the same species that share a freshwater adaptive zone (FAZ; Holtby and Ciruna 2007) or, if there were no such observations, from CUs in the closest FAZ that did have observations. Observations were weighted by the data quality score when calculating means (see Wilson and Peacock 2025). There were just two CUs that lacked any data on spawn timing (Boundary Bay Winter and Lower Fraser Summer steelhead), and in those cases we used spawn timing of nearby CUs of the same life-history type (Lower Fraser Winter and Fraser Canyon Summer, respectively).

For the adult freshwater migration stage of lake-type sockeye CUs, we considered the period of potential exposure starting with freshwater re-entry and extending for the duration of time it would take sockeye to reach their rearing lake, regardless of when spawning usually began. We based this migration duration on the distance from the ocean entry point to the rearing lake for the CU, with migration rates based on published data by run timing group (Quinn 2018; English et al. 2005; see Online Supplement for details). This accounts for the ability of lake-type sockeye CUs, particularly early-migrating CUs, to hold in lakes where they can moderate exposure to high temperatures by moving to deeper water and are not exposed to extreme flows.

We defined the spatial distribution of freshwater life stages by intersecting two spatial datasets: CU boundaries that encompass spawning watersheds used by each CU (Fisheries and Oceans Canada 2024) and modelled habitat suitability for spawning and rearing (Norris 2024). The estimated spawning and rearing suitability is an extension of previous work to model accessible fish habitat in BC accounting for both natural and anthropogenic barriers to migration (bcfishpass, Norris 2024). Spawning and rearing stream reaches were defined as a subset of accessible habitat based on species and life-stage-specific information related to mapped estimates of channel gradient, mean annual discharge, and minimum channel width (Table 1 of Rebellato et al. 2023). We assume that spawning and rearing stream reaches that fall within a CU boundary represent the available spawning and rearing distribution of that CU. One notable exception is the Harrison Upstream-Migrating lake-type sockeye CU (also known as Weaver Creek sockeye), for which rearing occurs upstream of the CU boundary in Harrison Lake, and this was accounted for when defining rearing stream reaches. The spatial extent of the adult freshwater migration stage was the river path between the mouth of the Fraser River and spawning stream reaches for the CU.

In the marine environment, our knowledge of both the timing and spatial distribution of life stages is less certain and less precise. There is compelling evidence that ocean conditions in the months preceding and around ocean entry at a regional scale influence early marine survival of salmon (e.g., Peterman et al. 1998; Mueter et al. 2002b, 2005). Therefore, we considered an early marine stage that occurred over a six-month period centered around the peak ocean entry date for each CU (Wilson and Peacock 2025). The spatial domain for the early marine stage extended 300 km south and 750 km north from the mouth of the Fraser River and 100 km offshore from the coastal shelf break, assuming that most CUs head north along the coast upon marine entry. Decades of studying the early marine survival of Fraser River salmon have yielded more detailed information on migration routes and timing of some CUs, particularly for Chinook (e.g., Trudel et al. 2009; Beamish 2018), but given that this information is not easily summarized for all CUs and the temporal and spatial resolution of available marine climate projections is relatively coarse, we opted for a simple and broadly applicable approach.

We assessed exposure for the marine rearing stage considering the remaining time in the life cycle from three months after ocean entry to the end of freshwater re-entry of returning adults. The spatial domain for marine rearing of all species encompassed the entire North Pacific Ocean and Bering Sea north of 35° N latitude. Although there have been efforts to define marine distribution at a species-level, these distributions are biased by unbalanced sampling and reliance on fisheries data and tend to vary with climate, particularly temperature, challenging their applicability under future climate scenarios (e.g., Langan et al. 2024).

We did not consider potential adaptation of life-history traits or spatial distribution in response to projected changes in climate variables. Currently, there is not sufficient data to characterize the relationships between climate variables and life-history timing or distribution of CUs. Life-history timing and distribution range were based on historical observations and treated as static under future climate scenarios. We consider the implications of this assumption on our results, and potential future research directions in this regard, in the Discussion.

For each life stage, we calculated exposure to relevant climate change variables (Table 2). The following sections describe the exposure calculations for each of these variables, separated by freshwater and marine habitats due to the different scales and data sources used to inform calculations in each. Finally, we describe how exposure was then integrated across habitats and life stages to yield an overall exposure for each CU.

**Table 2.**
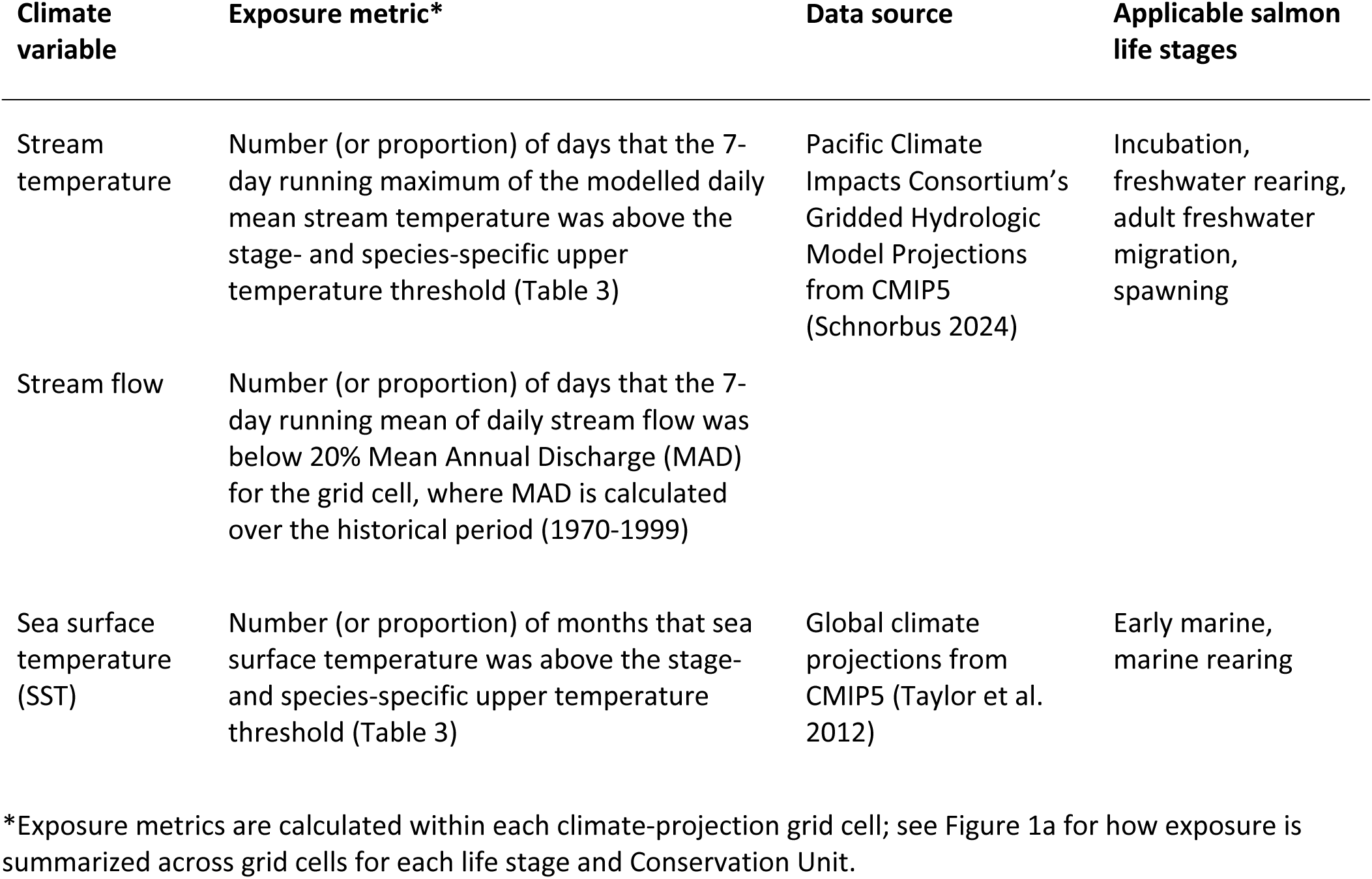
Overview of the climate variables, exposure metrics, and data sources used to quantify change exposure for each salmon life stage. See sections “Freshwater exposure” and “Marine exposure” for details.

| Climate variable | Exposure metric* | Data source | Applicable salmon life stages |
| --- | --- | --- | --- |
| Stream temperature | Number (or proportion) of days that the 7-day running maximum of the modelled daily mean stream temperature was above the stage- and species-specific upper temperature threshold (Table 3) | Pacific Climate Impacts Consortium’s Gridded Hydrologic Model Projections from CMIP5 (Schnorbus 2024) | Incubation, freshwater rearing, adult freshwater migration, spawning |
| Stream flow | Number (or proportion) of days that the 7-day running mean of daily stream flow was below 20% Mean Annual Discharge (MAD) for the grid cell, where MAD is calculated over the historical period (1970-1999) |  |  |
| Sea surface temperature (SST) | Number (or proportion) of months that sea surface temperature was above the stage- and species-specific upper temperature threshold (Table 3) | Global climate projections from CMIP5 (Taylor et al. 2012) | Early marine, marine rearing |

\*Exposure metrics are calculated within each climate-projection grid cell; see Figure 1a for how exposure is summarized across grid cells for each life stage and Conservation Unit.

### Freshwater exposure

Freshwater exposure was based on downscaled hydrologic model projections of daily mean stream temperature and flow; available from the Pacific Climate Impacts Consortium (PCIC; pacificclimate.org). PCIC’s gridded streamflow and water temperature data included projections for daily mean stream temperature and flow from 1945 to 2099, on a spatial grid with a resolution of 1/16th of a degree for both latitude and longitude (Schnorbus 2024). This output was available for six different Global Climate Models (GCMs) from the Coupled Model Intercomparison Project Phase 5 (CMIP5) ensemble, under the RCP 4.5 and RCP 8.5 representing moderate and high emissions scenarios, respectively (Taylor et al. 2012). For a description of the emission scenarios and list of the GCMs and specific simulations that we drew on, see the Online Supplement (Table S2). All exposure calculations described below were repeated for each of the six GCMs and two emissions scenarios.

#### Stream temperature

Stream temperature can directly affect salmon during incubation, juvenile rearing, adult freshwater migration, and spawning. For example, eggs have a narrow thermal tolerance and may experience thermal stress and oxygen limitation over summer or in early fall (Martin et al. 2017). At higher stream temperatures, enroute mortality of return-migrating salmon has been shown to increase in the Fraser River (Rand et al. 2006; Martins et al. 2011) and elsewhere (e.g., Crozier et al. 2020; Atlas et al. 2021). Stream temperature has been generally increasing, particularly over the summer months when the effects of increasing air temperatures are compounded by lower flows. Summer water temperatures in the Fraser River have warmed by ∼1.5 °C since the 1950s (Patterson et al. 2007). The variability in the current thermal regime of rivers and in projected climate changes throughout BC (Weller et al. 2023) mean that the magnitude of temperature increases experienced by salmon may differ among watersheds and CUs.

For each grid cell, we smoothed projected stream temperature using the 7-day running maximum of the modelled daily mean temperature. The 7-day mean of the daily maximum temperature is widely regarded as the most relevant metric for fish exposure to temperature (United States Environmental Protection Agency 2003), but projections of daily maxima were not available. Our results of exposure to stream temperatures are therefore likely conservative as the daily maximum may be ∼ 3°C above the daily mean temperature minimum (Oliver and Fidler 2001), but this bias is consistent among CUs and does not affect relative exposure rankings. For each grid cell that intersected the spatial distribution for a given life stage, we calculated the number of days within the stage window that the 7-day running maximum exceeded the temperature threshold (Table 3).

**Table 3.**
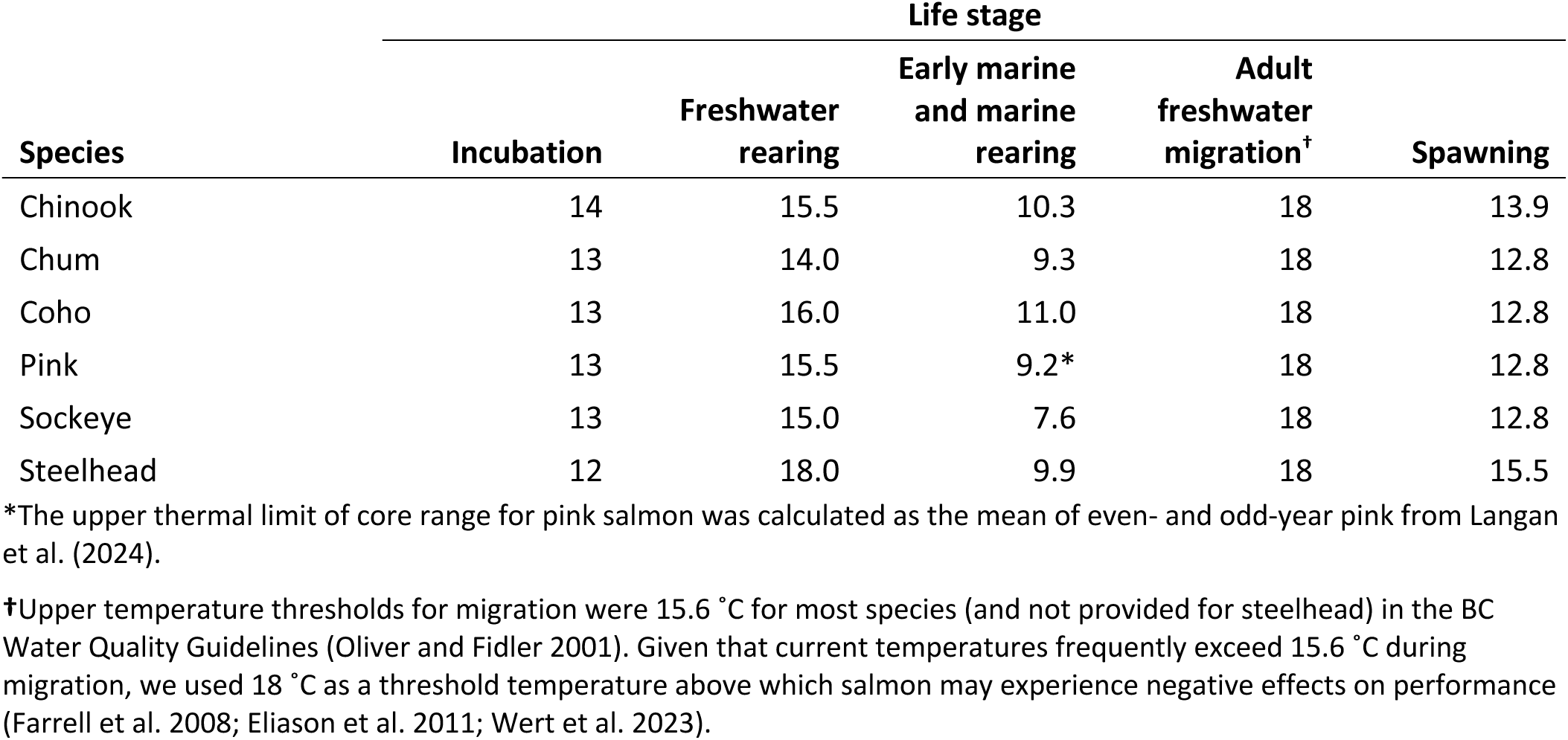
Upper temperature thresholds (*°*C) were chosen as temperatures above which there is evidence of adverse effects on a species as described in Oliver and Fidler (2001) or, for the marine stages, the upper temperature of their core thermal range (Langan et al. 2024).

Maximum temperature thresholds for salmon differ among species and life-stages (Richter and Kolmes 2005), and within species (see Discussion). We aimed to quantify salmon exposure to temperatures that may have sublethal adverse effects on behaviour and ecology of salmon, rather than temperatures that cause direct mortality (Mayer et al. 2023). We based upper temperature thresholds on species- and life-stage specific guidelines from the Province of BC (Oliver and Fidler 2001), which were developed through a literature review of studies examining the thermal tolerance and preference of salmonids. We adjusted the maximum threshold during adult migration to 18 °C for all species, as suggested by other reviews (e.g., Richter and Kolmes 2005), because the BC Guidelines were much lower than commonly observed temperatures for this stage. See the Online Supplement (Figures S1-S5) for a comparison of our assumed temperature thresholds to other studies reported by Mayer et al. (2023).

#### Stream flow

Climate change is impacting the hydrology of freshwater ecosystems by altering precipitation patterns, reducing the snowpack and hastening glacial retreat, and changing the timing of spring snow melt, leading to more variable flows and shifting the timing of peak flows (Islam et al. 2019). The impacts of changing hydrographs on salmon will depend on the season and life stage. We focused on exposure to low flows, which are more easily quantified using available data across watersheds and CUs than exposure to damaging high flows. Although high flows are an increasing concern for some Fraser River CUs (Patterson et al. 2023), the extent to which high flows adversely impact salmon will depend on the hydrogeomorphology of stream reaches and was difficult to quantify across our broad study region.

We smoothed projected flow using the 7-day running mean of the daily flow to better capture extended periods of low flow that may be harmful to fish. As for stream temperature, for each grid cell that intersected the spatial distribution for a given life stage, we calculated the number of days that flow was below the threshold. We chose a threshold of 20% mean annual discharge (MAD), which is commonly applied when managing environmental flow needs and a level below which indices of fish abundance or survival generally decline (Rosenfeld and Enright 2022). The 20% MAD threshold was calculated for each grid cell using the mean annual discharge in the historical period (i.e., mean of all daily flows in 1970-1999).

### Marine exposure

Projections of marine climate variables were provided by NOAA (Jamie Scott, pers. comm; NOAA 2024) for ocean variables including sea surface temperature (SST) and sea surface salinity. To match the GCMs and emissions scenarios available for the freshwater projections, we considered the same six GCMs from the CMIP5 ensemble under the RCP 4.5 and RCP 8.5 scenarios (see Online Supplement) (Taylor et al. 2012). The marine projections were aggregated as monthly averages for 1901-2100 with a spatial resolution of 1 degree longitude × 1 degree latitude (approximately 70 km × 110 km). We focused on quantifying exposure of marine stages to projected changes in SST but also consider exposure to sea surface salinity in the Online Supplement.

#### Sea surface temperature

Analyses of salmon spawner-recruit data have repeatedly demonstrated relationships between survival and SST across species and populations (e.g., Mueter et al. 2002b; Connors et al. 2020; Murdoch et al. 2024). The mechanism of impact is likely multi-faceted, with potential direct impacts on physiological performance and (more commonly) indirect impacts via changes to prey availability and quality, competitive interactions, and predation (Beauchamp et al. 2007). The direction and magnitude of the relationship between salmon marine survival and SST can differ with latitude and through time but for salmon in the Fraser River basin, there is compelling evidence that positive anomalies in SST are associated with reductions in survival across species (Welch et al. 2000; Mueter et al. 2002a; Sharma et al. 2013; Connors et al. 2020; Malick 2020).

To facilitate comparisons of exposure among freshwater and marine life stages, we take a similar approach as with stream temperature and stream flow and quantify the number of months that salmon are exposed to temperatures above an upper threshold during the early marine or marine rearing stages for each grid cell within the spatial domain for the stage. Upper temperature thresholds were based on the upper limit of the core thermal range for each species (Table 3Table *3*), as described by Langan et al. (2024) in their analysis of ocean catches of salmon and steelhead in the North Pacific from 1953-2022. This assumes that salmon are preferentially selecting regions in the marine environment that are thermally optimal for growth and survival and is not based on experimental evidence or observations of adverse effects above this threshold. These upper thermal limits are consistent with a previous analysis of thermal habitats of Pacific salmon (Abdul-Aziz et al. 2011). We apply these thresholds to both early marine and marine rearing stages, but SST is likely higher in nearshore waters and the early marine stage may be tolerant of SST above the threshold we consider.

### Summarizing exposure

We summarized exposure of CUs to climate change variables for each life stage (Table 2) across GCMs for different periods and emissions scenarios (Figure 1). To capture trends in exposure due to anthropogenic climate changes, and minimize the impacts of stochasticity in the models on results (Schoeman et al. 2023), we averaged the duration above or below a threshold for each climate variable (Table 2) over each of four 30-year periods: historical (1970-1999), early century (2010-2039), mid century (2040-2069), and late century (2070-2099).

**Figure 1.**
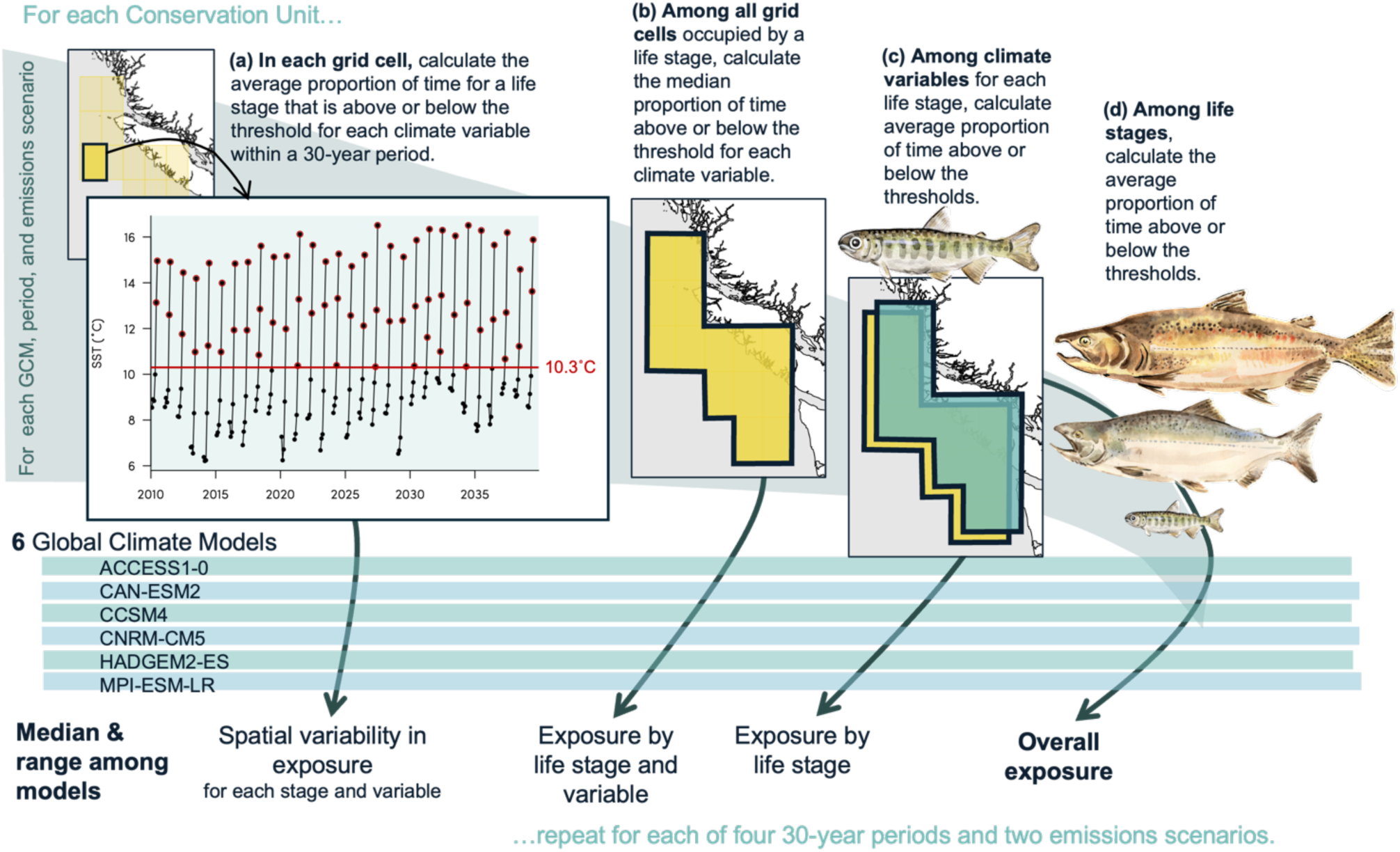
Illustration of our framework to assess climate change exposure of Pacific salmon and steelhead Conservation Units. (a) Exposure for each life-stage and climate variable (Table 2) was calculated as the average proportion of the stage duration that was above (or below) climate-variable thresholds. Illustrated here are SST projections for the early marine rearing stage of a Chinook CU in the early century period, with monthly temperatures above 10.3 °C highlighted by a red outline. In this case, there was an average of 2.3 out of 6 months above threshold, corresponding to a proportion (i.e., exposure) of 0.383 (b) The exposure of the early marine stage of this CU to SST was then calculated as the median proportion among all grid cells that overlapped the spatial distribution of the life stage. (c) The climate change exposure of the life stage was calculated from the relevant climate variables – for marine stages we considered SST only but for freshwater stages this was the average between stream temperature and low flow. (d) Finally, overall exposure was calculated as the average exposure among life stages (both freshwater and marine). At each of these steps, exposure is reported as the median and range among the six Global Climate Models (bottom). These calculations were repeated for four periods and two different emissions scenarios, for each of 60 Conservation Units.

Within each period, we summarized exposure for each CU and life stage by first calculating the number of days that the climate variable was above or below threshold (the specific metric depended on the variable; see Table 2). To standardize a metric of exposure among stages with different durations, we divided the numbers of days by the duration of the stage to yield the **proportion** of time above or below threshold (Figure 1a). If a stage was distributed over more than one grid cell, we summarized the exposure of the stage as the median proportion among all relevant grid cells (Figure 1b). To calculate overall exposure of freshwater life stages, we averaged exposure between stream temperature and flow; Figure 1c). Finally, to calculate overall exposure of the CU, we took the average of exposure among the six life stages (Figure 1d). This assumes an equal weight of each life stage to overall exposure. We also considered weighting exposure by the life-stage duration, but this led to exposure of the marine rearing stage tending to dominate outcomes. Future work could explore the impact of weighting stages differently based on, for example, their demonstrated impact on reproductive success.

For summaries of exposure within each variable and life stage (in each grid cell or among grid cells), across variables, and across life stages, we report the median and range in exposure values among GCMs, where the range represents uncertainty in climate model projections. This accounts for process uncertainty around how global climate changes will unfold, as captured by the different GCMs and their associated assumptions. Maintaining the distinct outcomes under different GCMs as we moved through each summary also allowed us to capture the greater range in overall exposure outcomes among GCMs that might be expected if exposure in marine and freshwater environments is correlated within models.

We focus on reporting results for the mid-century period, which is a future period that may be most relevant for developing and investing in meaningful strategies to improve salmon resilience to climate changes. Similarly, we focus on the RCP 4.5 emissions scenario (Burgess et al. 2023), and discuss the RCP 8.5 outcomes only when there were notable differences. Results for all periods and emissions scenarios are available in the Online Supplement.

## Results

### Species summaries

In general, overall exposure to climate changes across life stages was greatest for sockeye salmon (Figure 2, Figure 3Figure 3). All sockeye CUs had relatively high exposure to increasing SST during the early marine stage (Figure 4, Table S5), owing to the lower marine thermal preference of sockeye compared to other species (Table 3). However, sockeye also exhibited the greatest range in overall exposure among CUs due to the diverse timings of freshwater life stages among CUs. For many sockeye CUs, especially early-summer-migrating (ES) CUs, the spawning stage was highly exposed to stream temperatures above threshold due to the combination of a relatively low thermal tolerance, late-summer spawn timing (Table S1), and more constrained (often interior) distribution of CU-specific spawning habitats. Lake-type sockeye CUs that spawn later, including Cultus-Late, Seton-Late, Lillooet-Harrison-Late, and Shuswap-Late, generally had much lower exposure of the spawning stages to stream temperatures above threshold.

**Figure 2.**
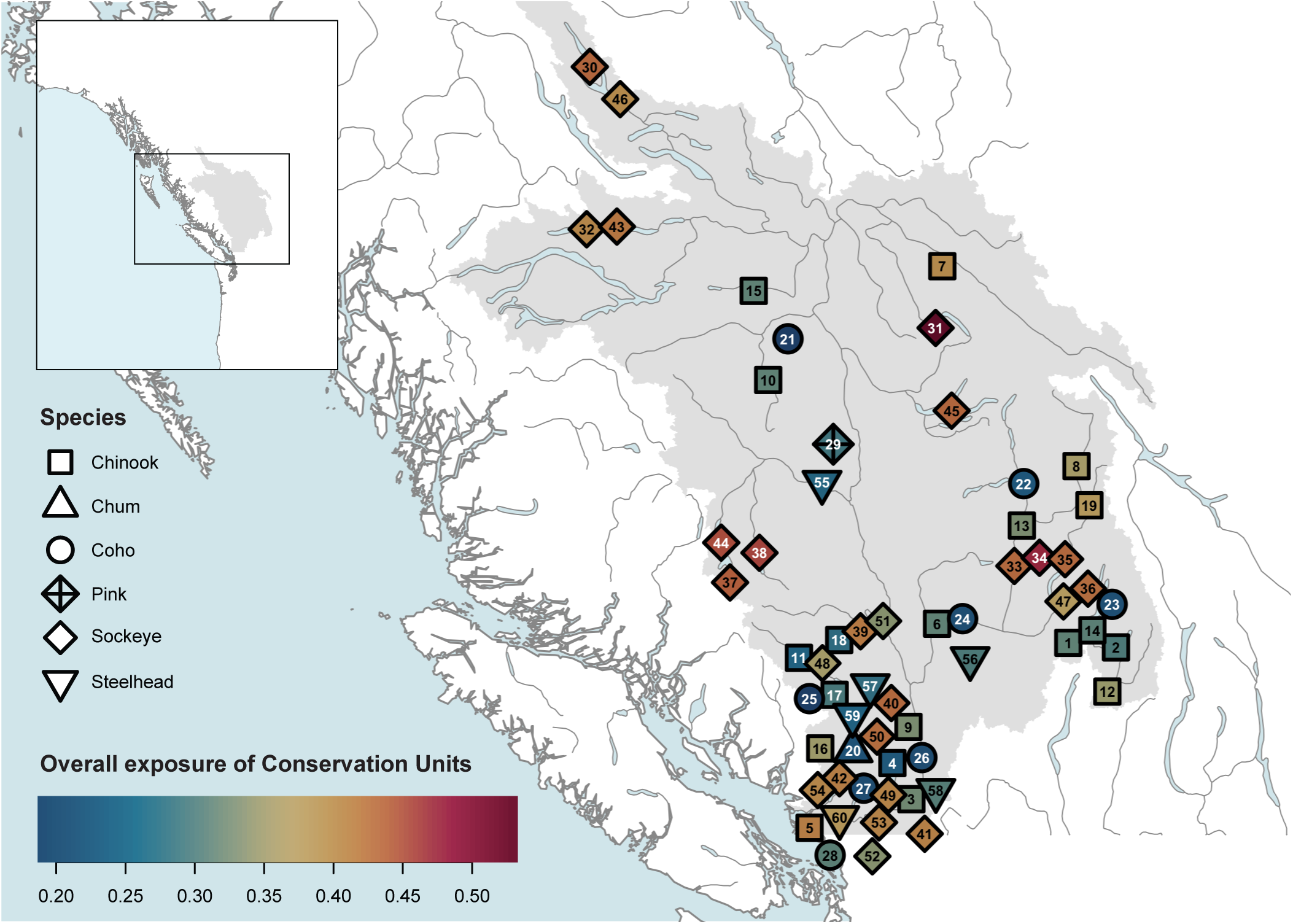
Map showing the Fraser River basin (grey) and the approximate locations of 60 Conservation Units (centroids of boundaries) for six species of Pacific salmon (point types; see legend on left). Points are coloured by their overall exposure to climate change for the **mid-century period (2040-2069) under the RCP 4.5 emissions scenario**. See Figure 3 and Table S1 for corresponding names of numbered Conservation Units on map.

**Figure 3.**
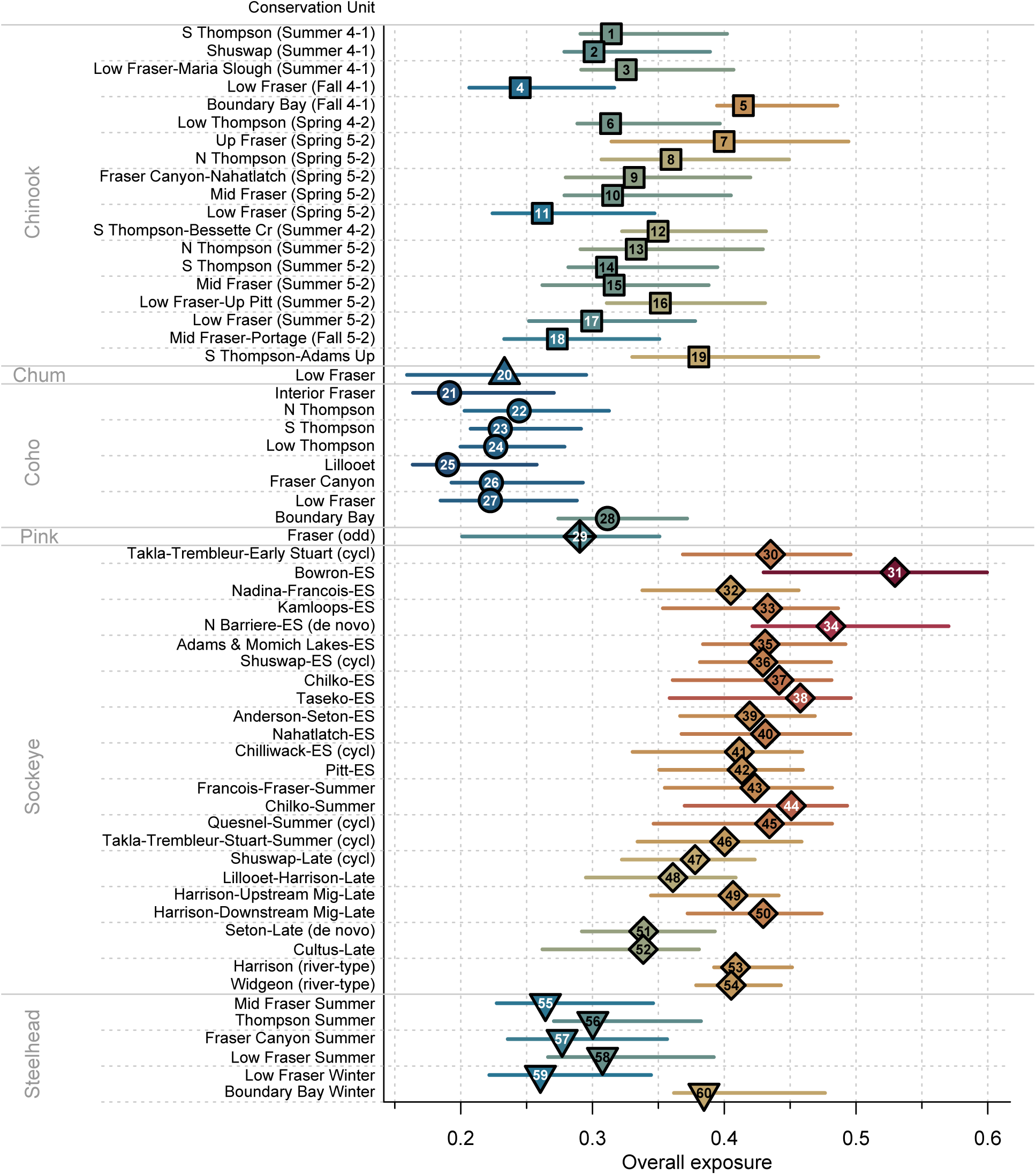
Overall exposure to climate changes across the lifecycle (x-axis) for 60 salmon CUs in the Fraser River basin (y-axis) for the **mid-century period (2040-2069) under the RCP 4.5 emissions scenario**. Points and lines are the median and range in exposure among six different GCMs. CUs are ordered by species and, within species, by life-history type and approximate distance from the river mouth (starting with the furthest migrating CUs), with numbers and shading corresponding to points in Figure 2. Exposure was calculated as the proportion of the lifecycle that was above or below the thresholds for different climate variables, averaged among variables and life stages for each CU. See Methods for details and Online Supplement for other periods and emissions scenarios.

**Figure 4.**
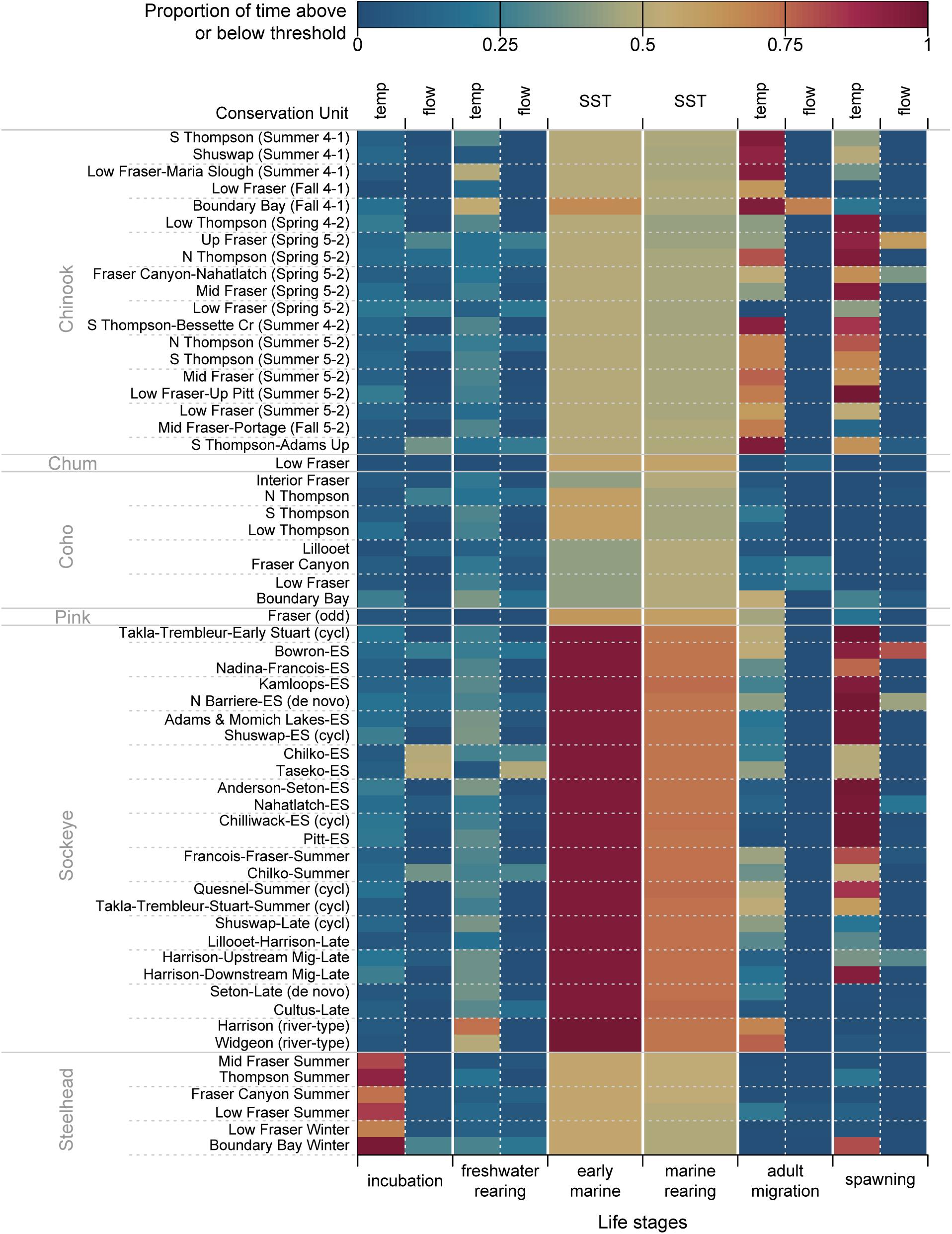
The proportion of time above threshold (for temperature) or below threshold (for flow and salinity) for each stage and climate variable (x -axis) and each CU (y-axis) in the **mid-century period (2040-2069) under the RCP 4.5 emissions scenario**. Stream temperature (temp) and low flow (flow) were assessed for freshwater stages; sea surface temperature (SST) was assessed for marine stages. CUs are ordered by species and, within species, by life-history type and approximate distance from the river mouth (starting with the furthest migrating CUs). See Methods for details and Online Supplement for other periods and emissions scenarios and exposure of marine stages to sea surface salinity.

The two river-type sockeye CUs, which do not rear in freshwater like their lake-type counterparts, had moderate overall exposure, and near-zero exposure of spawning stages to high stream temperature because they spawn later and in more coastal watersheds where there is expected to be less warming. However, these river-type CUs had relatively high exposure during the adult freshwater migration stage, which is distributed in the lower Fraser River in late July through early October when river temperatures are at their highest.

Chinook were generally the second most-exposed group of CUs. Overall exposure of Chinook CUs did not show obvious patterns among life-history types. Ocean-type (also known as 0+ or sub-yearling and denoted (4-1) in CU names to indicate an average generation length of four years and migration to sea in the first year of life) and stream-type (also known as 1+ or yearling and denoted (4-2) or (5-2)) had similar proportional time above temperature thresholds, but it is worth noting that the absolute duration that stream-type CUs might experience elevated stream temperatures would be longer. Although overall exposure was similar, the life stages that were driving exposure varied among migration timings, with Spring CUs having lowest exposure of the adult migration stage but higher exposure of the spawning stage. Conversely, Fall CUs had the highest exposure of the adult migration stage and lowest exposure of spawning stages.

The spatial distribution of Chinook CUs seemed to play more of a role in determining exposure. Interior CUs with relatively long migrations, including the Upper Fraser (Spring 5-2), South Thompson-Adams River Upper, and the North Thompson (Spring 5-2), were among the most exposed Chinook while the Lower Fraser (Fall 4-1) and Lower Fraser (Spring 5-2) were the least exposed. One exception to this pattern was the Boundary Bay (Fall 4-1) CU which has a small and very urban distribution in BC’s Lower Mainland and is projected to have especially high exposure to stream temperatures and low flows during the adult migration stage (as did the Boundary Bay Winter steelhead).

Steelhead had moderate overall exposure, but all six steelhead CUs had notably high exposure of the incubation stage to stream temperature. Steelhead spawn in the spring and therefore eggs incubate over the summer months (Table S1; Wilson and Peacock 2025), unlike other salmon, which tend to spawn in the fall or winter and incubate over winter. Further, steelhead eggs have the lowest thermal tolerance of the six species we considered (Table 3).

There was apparently low exposure of steelhead to high stream temperature during adult migration and holding, which can last almost a year for summer run populations. However, this was likely because the long duration of this stage results in a lower proportion of days above threshold, but the absolute number of days that steelhead experience stream temperatures over threshold would be similar to other CUs that inhabit the same streams over summer (see Table S5 for a comparison of days over threshold versus proportion of stage duration over threshold for each variable). Other stages had moderate to low exposure except for Boundary Bay Winter steelhead, for which the spawning stage was highly exposed to stream temperatures above threshold.

Coho salmon were among the least exposed CUs (Figure 3) despite spending at least a year in freshwater as juveniles. Coho juveniles often rear in small streams and side channel habitats and have thus evolved to be relatively tolerant to high temperatures (Table 3). Coho salmon also tend to spawn later than other species (Table S1), meaning that returning adults, spawners, and eggs do not experience the heat of late summer like other CUs.

Pink and chum are each considered a single CU in the Fraser River basin, and both had relatively low exposure. Pink and chum salmon enter the marine environment soon after fry emergence; therefore, the exposure of the freshwater juvenile stage was zero in our assessment. Pink and chum had slightly higher exposure than most coho and Chinook CUs in the early marine stage (Figure 4) because they had lower core marine temperature ranges (Table 3).

### Climate variables and stages driving exposure

Exposure to stream temperatures above the threshold during the freshwater adult migration and spawning stages tended to differentiate CUs more than other climate variables or stages (Figure 4). For freshwater adult migration and spawning, both spatial distribution and timing varied considerably among CUs, leading to varying degrees of exposure (e.g., Figure 5).

**Figure 5.**
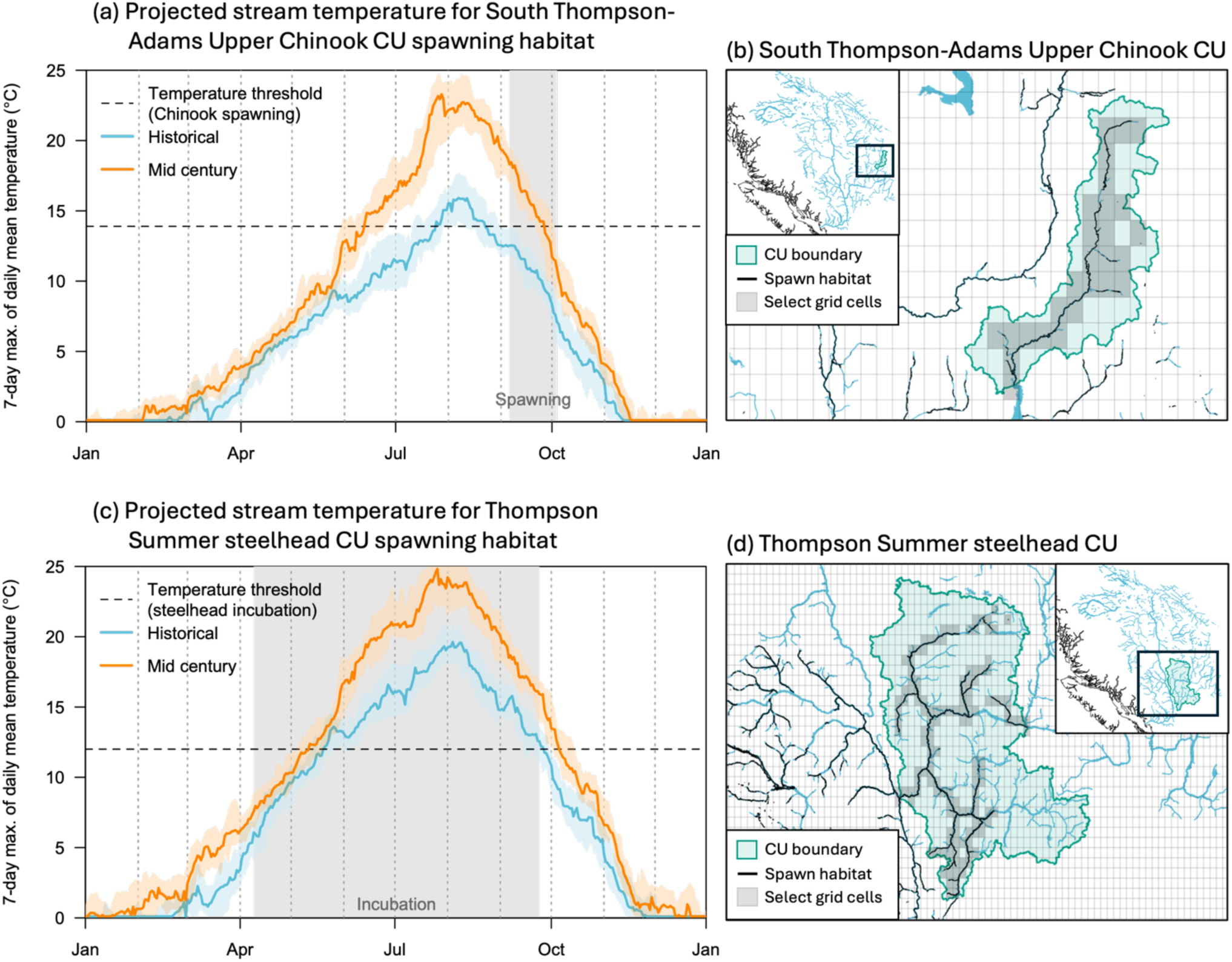
Examples of exposure to stream temperatures in the mid-century period (2040-2069; orange) relative to the historical period (1970-1999; blue) for two CUs: (a-b) The South Thompson-Adams River Upper Chinook CU had increasing exposure of the spawning stage to elevated stream temperatures by the mid-century period. (c-d) The Thompson Summer steelhead CU had relatively low overall exposure, but the incubation stage was notably exposed to stream temperatures above the threshold. Maps in (b and d) show the locations of these CUs with the 1/16° grid of hydrologic projections overlaid. Select grid cells relevant to the spawning and incubation stages, over which the temperatures in (a) and (c) were summarized, are shaded grey and overlapped modelled spawn habitat for the CU. The inset maps show the location of the CU within the Fraser River basin. Stream temperature projections shown are from the Can-ESM2 GCM under RCP 4.5.

Exposure to low flows was minimal for most CUs, with some notable exceptions. For CUs that spawn in the interior of the Fraser River basin, including the Bowron-ES lake-type sockeye and Upper Fraser Spring (5-2) Chinook, flows were predicted to decrease during spawning periods, leading to relatively high exposure of that stage to low flows (Figure 6). However, for these same CUs, flows may increase during other times of the year and lead to reduced exposure of other stages to low flows. For example, flows in the interior northeast of the Fraser River basin are generally projected to increase over winter as precipitation increases, leading to lower exposure of incubation and rearing stages to low flows (e.g., Figure 6a). Thus, the exposure of CUs to low flows in future periods depended both on the projected shifts in timing and duration of annual peak flows and their life-cycle timing.

**Figure 6.**
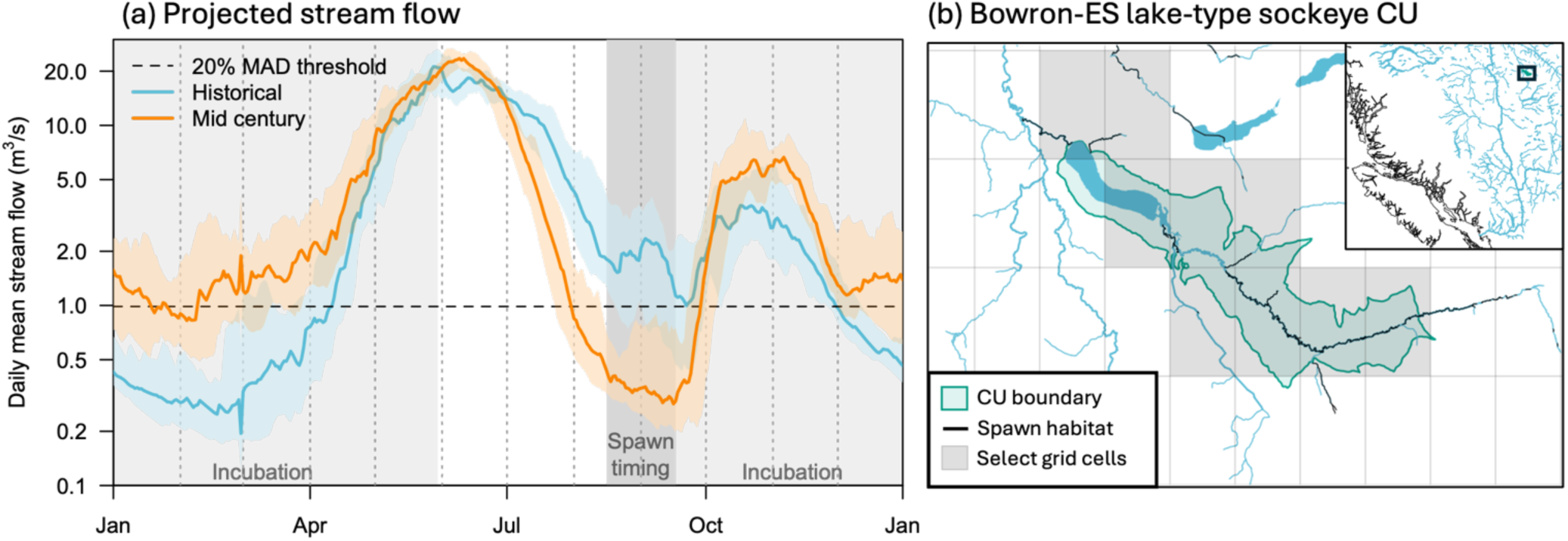
(a) Example of increased exposure of the spawning stage to low flows below 20% MAD for the Bowron-ES lake-type sockeye CU in the mid-century period (2040-2069; orange) relative to the historical period (1970-1999; blue). Flows are shown as the median (solid line) and interquartile range (semi-transparent envelope) of daily mean flows among 30 years within each period from the Can-ESM2 GCM under RCP 4.5. Exposure to low flows during the incubation period, however, declined from the historical period to the mid-century period. (b) Map of the Bowron-ES lake-type sockeye CU with the 1/16° grid of hydrologic projections overlaid. Select grid cells relevant to the spawning and incubation stages, over which the flows in (a) were summarized, are shaded grey and overlapped modelled spawn habitat for this CU. The inset map shows the location of the CU, in the northeast of the Fraser River basin.

For marine stages, CUs were assumed to have similar timing, the same broad spatial distributions, and similar exposure thresholds so there was less differentiation in exposure among them at this life stage. Differences in exposure of marine stages were mostly due to the assumed species-specific core thermal range. Though not included in our overall exposure, we also note that there was almost no difference among CUs in exposure to changing salinity, because the thresholds were based on historical distribution of values and were the same for all species and CUs (Online Supplement).

### Patterns over time periods and emissions scenarios

In general, exposure tended to increase from early to mid century but then plateau, or even decline, in the late century under both the low and high emissions scenario (Online Supplement). The decline in exposure towards the late century was not observed for all CUs and stages. Some CUs, however, saw a decline in exposure to low flows and high stream temperatures during the adult freshwater migration and spawning stages. There was considerable disagreement among GCMs about declines in exposure into the late century, highlighting the model uncertainty around long-term projections and warrant caution when interpreting changes into the late century.

## Discussion

Our assessments highlighted how life-history timing and stage-specific habitat use of different salmon and steelhead CUs can influence their relative exposure to climate changes. We found that sockeye generally had the highest exposure to climate changes throughout their lifecycles but also showed the highest diversity of exposure outcomes among CUs owing mainly to different migration and spawn timings. Coho, pink, and chum salmon had the lowest overall exposure due to both their relatively late migration and spawn timings and (in the case of pink and chum salmon) short duration of freshwater stages and higher thermal tolerance. Our findings align with previous assessments of salmon climate exposure in the Pacific Northwest (e.g. Wade et al. 2013; Crozier et al. 2019), but CU-specific nuances offer new insight into how climate change may affect diverse Pacific salmon populations in Canada.

### Adaptive capacity

Moving from our estimates of exposure to assessing climate change vulnerability, which is defined as “the degree to which a system is susceptible to, and unable to cope with, the adverse effects of climate change” (Schneider et al. 2007), requires understanding the adaptive capacity of the CUs to change in ways that enable survival in changing conditions. We found that exposure outcomes were highly dependent on salmon life-history traits, which emphasizes the need to better understand how salmon may adapt their life cycles under climate change.

We treated CU life-history timing as static (Wilson and Peacock 2025), yet inter-annual variation shows salmon can adjust to current environmental conditions. Broad spatial patterns in freshwater life histories also indicate adaptive capacity - for example, Chinook runs occur in spring and fall in southern watersheds but converge to summer at more northern latitudes with lower stream temperatures (Potapova et al. 2026). Modelling suggests sockeye migration timing can adapt quickly enough to reduce thermal exposure in the Fraser River (Reed et al. 2011). Despite this apparent plasticity in life-history traits, shifts in phenology may involve trade-offs affecting the growth and survival of later stages (Crozier et al. 2008).

Adaptation is not limited to altering life-history strategies; it can also involve tolerating environmental changes. While we assumed that thermal ranges were similar among CUs, evidence shows local thermal adaptation within salmon species (e.g., Eliason et al. 2011; Mayer et al. 2023). Over the intergenerational timescales relevant to climate change projections, CUs may retain some capacity to expand their thermal tolerance. However, climate change could also push cold-water fishes to the edge of their physiological limits (Adams et al. 2022), constraining this form of adaptation (Muñoz et al. 2015). Similarly, we applied a common 20% MAD low-flow threshold for all CUs and life stages, yet research suggests that more conservative thresholds may be necessary for certain stages (Bradford et al. 2011). For example, Warkentin et al. (2022) found that Chinook salmon in the Nicola River watershed would require 36% MAD during rearing to support a stable population. These results point to the importance of developing stage-specific environmental guidelines and improving our understanding of the scope for adaptive capacity through tolerance.

The spatial distribution of CUs was another key factor differentiating exposure of freshwater stages, and there were notable uncertainties in the freshwater rearing stage that may influence results. In particular, the distribution of freshwater rearing for species like Chinook and coho was likely underestimated, especially for CUs that spawn in the upper Fraser River. We considered freshwater rearing to occur within the CU boundary, but salmon may migrate downstream to other watersheds during their freshwater rearing stage (Bradford et al. 2008; Bradford and Taylor 2023). This ability to move and seek out suitable habitats in other watersheds may reduce the potential exposure of the freshwater rearing stage to climate changes, and represents another form of adaptation that needs to be considered in assessing climate change vulnerability.

We could not resolve differences in exposure of marine stages among CUs, due to both a lack of CU-specific information on sensitivity thresholds and spatial distribution. We focused on quantifying exposure of marine stages to SST (and consider salinity in the Online Supplement) but many other marine climate and ecosystem variables would likely be relevant indicators for salmon (e.g., sea surface pH, oxygen, primary productivity). However, without clear relationships between those variables and salmon responses, it is difficult to meaningfully assess their potential impact. The similar exposure to decreasing salinity among CUs in this study (Online Supplement) demonstrated that, even if we know a climate variable is important, it may not be useful to understanding relative climate change exposure without information on species- or population-specific thresholds of impact. This challenge was also noted by Crozier et al. (2019) in their climate change vulnerability assessments, which found similar exposure among salmon populations to ocean acidification. Further research is needed to understand the responses of different salmon populations to changing ocean conditions, along with more comprehensive sampling to expand our understanding of CU-specific marine migration patterns (e.g., Weitkamp 2010; Freshwater et al. 2021).

### Refining climate projections

Higher resolution climate model projections will help further differentiate exposure of CUs in both marine and freshwater environments. Downscaled marine climate projections have been undertaken for the BC coast drawing on a single GCM (Holdsworth et al. 2021; Peña and Fine 2023) but the available output is limited to summaries within multi-decade future periods. As the storing and sharing of large datasets gets easier and less expensive, preserving raw outputs across multiple models would facilitate application in assessments tailored to different study systems (e.g., across marine/freshwater divide as in this work) and improve the ability to track uncertainties. Further, encouraging international cooperation to facilitate modelling across international borders would greatly improve the ability to assess climate change exposure of migratory and wide-ranging species, including salmon (Ferriss et al. In press.). Gaps in hydrologic model output for transboundary watersheds introduced some uncertainty in exposure of freshwater stages for small CUs distributed along the southern Canadian border (e.g., Boundary Bay Chinook, coho, and steelhead).

In contrast to the coarse marine projections, the hydrologic model projections we relied on for freshwater exposure were available at a finer daily temporal resolution and a finer spatial grid of 1/16 degree, but nonetheless this gridded output was an abstraction of the actual conditions experienced by fish. Stream temperature and flow projections within grid cells will tend to reflect the largest order streams within that grid cell, but smaller streams and side channels are important habitats for salmon that may provide climate refuges – or be lost under more frequent drought conditions. Model projections of daily mean temperatures and flows are likely to underestimate the extremes that salmon and other aquatic species experience, and underestimate the true exposure of aquatic species to climate change. Channel-level statistical models of August mean temperature (Weller et al. 2023) provide the opportunity to examine finer-scale changes within salmon habitats that would complement this analysis. Channel-level projections of stream flow would also allow for assessing exposure to high flows, which we were not able to do with the gridded output. The distinction between beneficial high flows, which are needed to remove sediment and maintain spawning gravels, and detrimental high flows, such as a 100-year flood event that will flush out eggs and juvenile salmon or inhibit adult migration, will depend on the specific hydromorphology of a river. Ongoing research will no doubt improve the quality and resolution of hydrologic projections that can be used to refine assessments of climate change exposure.

Another key limitation of the hydrologic modelling we used is the current lack of consideration given to how landscape disturbances will affect stream temperature and flow. Incorporating information on landscape disturbances is difficult as we cannot know for certain the extent of wildfires, logging, or road development (for example) decades into the future. But there is ample evidence that these disturbances contribute to more extreme flows and higher stream temperatures (e.g., Naman et al. 2024), potentially exacerbating the impacts of climate change on habitat suitability for salmon. Indeed, modelling studies suggest that landcover change is an important factor mediating future hydrologic conditions under climate change (Smith et al. 2023). Therefore, it is important that the status of freshwater habitats be considered alongside these climate change exposure outcomes when assessing risks to salmon viability and developing conservation strategies (Moore et al. 2025).

There are many other direct and indirect impacts of climate change or mediating factors that we could not address here, leaving ample room for improvement and additions in future work. For example, recent research has demonstrated that lake depth is an important predictor of sockeye responses to rising temperatures (Price et al. 2024), and yet the hydrologic models that we relied on to quantify exposure to high stream temperature included a relatively simple 2-layer lake model (Schnorbus 2024) that may not capture the variability among nursery lakes in their response to climate change (e.g., Putt et al. 2019). Similarly, there is increasing recognition that thermal refugia and small-scale heterogeneity in stream temperatures resulting from groundwater influx allow salmon to seek out thermally suitable habitat, and may be key to supporting diverse life-history types under future climate change (Snyder et al. 2022). The ocean environment displays similar nuances, and capturing the impact of these is further complicated by the flexibility of migration routes and residence times in the marine environment. These complexities may be impossible to incorporate directly into models but should be taken into consideration when interpreting studies like ours. More complex and cascading impacts of climate change on freshwater and oceanic food webs that support salmon are already being observed (Mackas et al. 2007), but our ability to predict and quantify potential impacts to salmon is currently limited.

## Conclusions

Information on climate change exposure is critical for salmon management in an era of rapid environmental change. The current salmon management paradigm remains largely focused on developing plans for rebuilding the most depressed stocks, but integrating information on climate change exposure alongside current habitat status and biological status enables a more forward-looking approach. Identifying CUs with lower exposure can guide protection of salmon strongholds (Wild Salmon Center 2021), while anticipating future stressors helps prioritize and future-proof recovery investments. As direct control of climate change challenges becomes increasingly unattainable and international cooperation fails, domestic and regional management must pivot towards mitigating impacts by fostering diverse and resilient salmon habitats (Moore and Schindler 2022; Moore et al. 2025). This aligns with strategic management frameworks such as RAD (Resist, Accept, Direct) (Williams and Brown 2024) or RRT (Resistance–Resilience–Transformation) (Peterson St-Laurent et al. 2021) which call for conservation strategies that evolve with accelerating change. The application and extension of these climate change exposure assessments can inform proactive protection and restoration that will deliver the greatest benefits among salmon and steelhead in Canada.

## Acknowledgements

We acknowledge the World Climate Research Programme’s Working Group on Coupled Modelling, which is responsible for CMIP, and we thank the climate modeling groups (listed in Table S2 of this paper) for producing and making available their model output. For CMIP the U.S. Department of Energy’s Program for Climate Model Diagnosis and Intercomparison provides coordinating support and led development of software infrastructure in partnership with the Global Organization for Earth System Science Portals.

We thank Simon Norris for providing output from the bcfishpass model that was used to define spawning, rearing, and migration habitats for each CU. We thank Travis Tai and Diana Dobson for early input on this work through PSF’s Climate Science Advisory Committee, and Jan Finke for comments on the final draft. We also thank members of PSF’s Population Science Advisory Committee and Habitat Science Advisory Committee for their feedback. This research was funded by the British Columbia Salmon Restoration and Innovation Fund, project #2020_27 led by the Pacific Salmon Foundation.

## Competing interests

We declare that we have no competing interests.

## Data availability

All code for these analyses are available at https://github.com/salmonwatersheds/ccva, and will be archived in Zenodo upon acceptance. Input data are publicly available via cited sources or included in the Online Supplement submitted with the manuscript (also available at https://salmonwatersheds.github.io/ccva/).

## Notes

https://salmonwatersheds.github.io/ccva/

### Competing Interest Statement

The authors have declared no competing interest.

### Summary of Updates

Revision to remove sea surface salinity results from the main text and address other comments from two reviewers; Figure 4 corrected and colours revised to emphasize climate exposure results rather than species.

