## Supplementary material for "Quantifying Exposure of Pacific Salmon and Steelhead to Climate Change in the Fraser River basin": Online Supplement: cjfas-2026-0084suppla.html

###### Stephanie Peacock

###### October 03, 2026

### Online Supplement

### 1 Conservation Units and their life-history traits

**Table S1.** List of the 60 Conservation Units in the Fraser River basin that were included in the study, along with life-history traits that were used to assess their exposure to climate changes.

Download as CSV

- Lake-type sockeye duration of exposure for adult freshwater migration was assumed to end when fish reached the rearing lake, where they are no longer exposed to potential low flows or high stream temperatures. See below for details.
  \*\* Adams & Momich Lakes-Early Summer (25 on map) is now considered two separate CUs: Adams - Early Summer (Extinct) and Momich - Early Summer (Endangered).

### 2 Global Climate Models

We acknowledge the World Climate Research Programme’s Working Group on Coupled Modelling, which is responsible for CMIP, and we thank the climate modeling groups (Table S2, below) for producing and making available their model output. For CMIP the U.S. Department of Energy’s Program for Climate Model Diagnosis and Intercomparison provides coordinating support and led development of software infrastructure in partnership with the Global Organization for Earth System Science Portals.

**Table S2.** The six Global Climate Models (GCMs) from which we used output in this paper and the research centres responsible for developing and running each GCM.

### 3 Temperature thresholds for freshwater stages

There is increasing evidence that salmonids can display considerable intraspecific diversity in thermal tolerance, likely due to local adaptation of populations to their freshwater thermal environments (Eliason et al. 2011; Zillig et al. 2021). This variability makes it difficult to determine an upper stream temperature threshold to quantify the future duration over threshold for freshwater life stages. Data on specific thermal tolerance of Conservation Units (CUs) is limited to a handful of CUs and life stages, limiting the applicability of this specific information in our broad-scale analysis. Further, a recent review highlighted the variability in outcomes under different experimental approaches, and the questionable ecological relevance of some of the reported metrics (Mayer et al. 2023). As such, we decided to rely on temperature guidelines set out by the Province of BC (Oliver and Fidler 2001) that specify a maximum optimal temperature for each species and stage, ignoring potential population-specific deviances.

To ensure the temperature thresholds we applied were not inconsistent with more recent research, we compared the BC Guidelines to the various studies reported on by Mayer et al. (2023) (Tables S2-S6 therein). These studies varied in their ecological relevance critical thermal maximum (CTmax) or upper incipient lethal temperature (UILT), which are considered to have low ecological relevance, to upper temperatures under which salmon can carry out activities essential to survival at any given life-stage (Pejus temperatures), considered to have high ecological relevance. We also show temperatures that are reported to be optimal or the upper observed range of populations in the wild.

There was considerable variability in thermal tolerance reported among studies, due to either population-specific differences or study methodology or both, but the BC Guidelines were not out of range of the thermal tolerance metrics reported by Mayer et al. (2023) (Figures S1 – S5). In general, the BC Guidelines were near upper-optimal or Pejus temepratures, which were considered the most ecologically relevant temperature thresholds by Mayer et al. (2023). The BC Guidelines were far below critical temperatures shown to cause direct mortality. This is consistent with our use and interpretation of the BC Guidelines: we aimed to quantify salmon exposure to temperatures that may have more subtle (sublethal) adverse effects.
Mayer et al. (2023) did not report any studies of thermal tolerance for spawning stages or for any life stage of steelhead that could be compared to the BC Guidelines.

Figure 3.1: Comparison of threshold stream temperatures for salmonid life stages from the BC Guidelines (Oliver and Fidler 2001) to more recently reported estimates of thermal tolerance summarized in Mayer et al. (2023). These other studies are categorized according to the ecological relevance of the approach (see legend and Mayer et al. 2023 for details).

Figure 3.2: Comparison of threshold stream temperatures for salmonid life stages from the BC Guidelines (Oliver and Fidler 2001) to more recently reported estimates of thermal tolerance summarized in Mayer et al. (2023). These other studies are categorized according to the ecological relevance of the approach (see legend and Mayer et al. 2023 for details).

Figure 3.3: Comparison of threshold stream temperatures for salmonid life stages from the BC Guidelines (Oliver and Fidler 2001) to more recently reported estimates of thermal tolerance summarized in Mayer et al. (2023). These other studies are categorized according to the ecological relevance of the approach (see legend and Mayer et al. 2023 for details).

Figure 3.4: Comparison of threshold stream temperatures for salmonid life stages from the BC Guidelines (Oliver and Fidler 2001) to more recently reported estimates of thermal tolerance summarized in Mayer et al. (2023). These other studies are categorized according to the ecological relevance of the approach (see legend and Mayer et al. 2023 for details).

Figure 3.5: Comparison of threshold stream temperatures for salmonid life stages from the BC Guidelines (Oliver and Fidler 2001) to more recently reported estimates of thermal tolerance summarized in Mayer et al. (2023). These other studies are categorized according to the ecological relevance of the approach (see legend and Mayer et al. 2023 for details).

Figure 3.6: Comparison of threshold stream temperatures for salmonid life stages from the BC Guidelines (Oliver and Fidler 2001) to more recently reported estimates of thermal tolerance summarized in Mayer et al. (2023). These other studies are categorized according to the ecological relevance of the approach (see legend and Mayer et al. 2023 for details).

### 4 Migration speeds and distances for lake-type sockeye CUs

The adult freshwater migration stage generally began with freshwater re-entry and extended until the start of spawning. For the adult freshwater migration stage of lake-type sockeye CUs, we considered the period of potential exposure extending for the duration of time it would take sockeye to reach their rearing lake, regardless of when spawning usually began. We based this migration duration on the distance from the ocean entry point to the rearing lake for the CU, with migration rates based on published data by run timing group (English et al. 2005; Quinn 2018). This accounts for the ability of lake-type sockeye CUs, particularly early-migrating CUs, to hold in lakes where they can moderate exposure to high temperatures by moving to deeper water and are not exposed to low flows. The following tables detail the data and sources used to estimate the duration of the adult freshwater migration stage for lake-type sockeye CUs.

**Table S3.** Data sources and values used to calculate the migration duration of lake-type sockeye CUs in the Fraser.

| Stock run timing group | Migration rate (km/day) | Sample size | Reference |
| --- | --- | --- | --- |
| Summer | 43 | 101 | English et al. (2005) Table 5 |
| Late | 20 | 96 | English et al. (2005) Table 5 |
| Early Stuart | 45.8 | NA | Quinn (2018) Table 4.3 |
| Horsefly (ES) | 51.2 | NA | Quinn (2018) Table 4.4 |
| Bowron (ES) | 51.5 | NA | Quinn (2018) Table 4.5 |

\***Table S4.** Lake-type sockeye CUs and associated start of freshwater re-entry, run timing group, migration rate (based on run timing group as outlined in Table S3 above), migration distance (calculated from rearing lake to point of ocean entry), and resulting duration of potential exposure.

Download as CSV

### 5 Supplemental results

#### 5.1 Sea surface salinity

In the main text, we consider exposure of marine life stages to sea surface temperature. Sea surface salinity is another oceanic variable that we considered for inclusion in the exposure assessments. Sea surface salinity can influence stratification and thus nutrient cycling in the ocean, driving bottom-up effects on salmon. Salinity in BC coastal waters has been decreasing since the middle of the last century (Chandler 2022) and long term projections of increased precipitation (Morrison et al. 2014) suggest that this trend will continue. There is some evidence of relationships between salinity and salmon growth or survival Mueter et al. (2005), but no information on thresholds or optimal salinity ranges.

Here, we present supplemental results that include the exposure of marine stages to sea surface salinities lower than the historical range that salmon have experienced. We focus on salinities *below* threshold under the assumption that projected decreases in surface salinity indicate increased stratification and reduced nutrient resupply to surface waters with general negative consequences for salmon (e.g., Thomson et al. 2012). We quantified the number of months below a lower salinity threshold, which we defined for each grid cell as the 2.5th percentile of the modelled historical (1970-1999) salinity. We note, however, that there is some uncertainty about how marine rearing stages will respond, and less saline conditions may have a net benefit for salmon in the open ocean.

In general, exposure to decreasing sea surface salinity was both low and consistent among CUs, offering little information to differentiate overall exposure or prioritise efforts to mediate climate-change impacts on salmon. As such, we opted to exclude these results from the main text, but highlight that more research is needed to understand how to relate proxies of oceanographic change, like sea surface temperature, to salmon survival.

Figure 5.1: Overall exposure to climate changes across the lifecycle (x-axis) for 60 salmon CUs in the Fraser River basin (y-axis) for the mid-century period (2040-2069) under the RCP 4.5 emissions scenario. Points and lines are the median and range in exposure among six different GCMs. Each CU has two version of exposure shown: semi-transparent points are the exposure *including* sea surface salinity for marine stages, and solid points are considering only SST for marine stages (as shwown in the main text). CUs are ordered by species and, within species, by life-history type and approximate distance from the river mouth (starting with the furthest migrating CUs), with numbers and shading corresponding to points in Figure 2. Exposure was calculated as the proportion of the lifecycle that was above or below the thresholds for different climate variables, averaged among variables and life stages for each CU. See below for other periods and emissions scenarios.

#### 5.2 Exposure outcomes

**Table S5.** The following table contains raw exposure outcomes (median and range among six GCMs) for each CU, life stage, and climate change variable (including sea surface temperature) under RCP 4.5 and 8.5 in the historical, early, mid, and late century periods. Exposure is presented as the proportion of stage duration over threshold for each variable in the main text, but also provided here as the absolute number of days over threshold for comparison.

##### Table

Download as CSV

##### Attributes

| Field | Description |
| --- | --- |
| **species** | CK = Chinook, CM = chum, CO = coho, PKO = odd-year pink, SEL = lake-type sockeye, SER = river-type sockeye, SH = steelhead |
| **cu\_name** | Name of the Conservation Unit |
| **cuid** | Unique identifier assigned by the Pacific Salmon Foundation to each Conservation Unit; can be cross-referenced with other PSF databases available from the Salmon Watersheds Data Library. |
| **stage** | One of six life stages distingished in the paper: `eggs_alevin`, `fw_rearing`, `early_marine`, `marine_rearing`, `adult_migration`, `spawning`. |
| **variable** | Climate change variable; for freshwater stages this is either `stream_flow` or `stream_temp` and for marine stages this is either `SSS` (sea surface salinity) or `SST` (sea surface temperature). |
| **rcp** | Moderate emissions (`rcp45`) or high emissions (`rcp85`) scenario. |
| **period** | One of four time perods: `hist` (1970-1999), `early` (2010-2039), `mid` (2040-2069), or `late` (2070-2099). |
| **exposure\_prop\_median** | median exposure among six GCMs, summarized as the proportion of the stage duration above threshold (for `stream_temp` and `SST`) or below threshold (for `stream_flow` and `SSS`). |
| **exposure\_prop\_min** and **exposure\_prop\_max** | The minimum or maximum exposure among six GCMs, summarized as the proportion of the stage duration above threshold (for `stream_temp` and `SST`) or below threshold (for `stream_flow` and `SSS`). |
| **stage\_duration\_days** | The duration of the life stage or the given CU in days. |
| **exposure\_days\_median** | Median exposure among six GCMs, summarized as the number of days above threshold (for `stream_temp` and `SST`) or below threshold (for `stream_flow` and `SSS`). Equal to the `stage_duration_days * exposure_prop_median`. |
| **exposure\_days\_min** and **exposure\_days\_max** | The minimum or maximum exposure among six GCMs, summarized as the number of days. |

#### 5.3 Overall exposure

##### Low emissions (RCP 4.5)

###### Historical

Figure 5.2: Overall exposure to climate changes across the lifecycle (x-axis) for 60 salmon CUs in the Fraser River basin (y-axis). The mid-century period (2040-2069) under the RCP 4.5 emissions scenario is included in the main text; **other time periods and emissions scenarios can be viewed here under the tabs.** Points and lines are the median and range in exposure among six different GCMs. CUs are ordered from highest (top) to lowest (bottom) overall exposure. Exposure was calculated as the proportion of the lifecycle that was above or below the thresholds for different climate variables, averaged among variables and life stages for each CU.

###### Early century

Figure 5.3: Overall exposure to climate changes across the lifecycle (x-axis) for 60 salmon CUs in the Fraser River basin (y-axis). The mid-century period (2040-2069) under the RCP 4.5 emissions scenario is included in the main text; **other time periods and emissions scenarios can be viewed here under the tabs.** Points and lines are the median and range in exposure among six different GCMs. CUs are ordered from highest (top) to lowest (bottom) overall exposure. Exposure was calculated as the proportion of the lifecycle that was above or below the thresholds for different climate variables, averaged among variables and life stages for each CU.

###### Mid century

Figure 5.4: Overall exposure to climate changes across the lifecycle (x-axis) for 60 salmon CUs in the Fraser River basin (y-axis). The mid-century period (2040-2069) under the RCP 4.5 emissions scenario is included in the main text; **other time periods and emissions scenarios can be viewed here under the tabs.** Points and lines are the median and range in exposure among six different GCMs. CUs are ordered from highest (top) to lowest (bottom) overall exposure. Exposure was calculated as the proportion of the lifecycle that was above or below the thresholds for different climate variables, averaged among variables and life stages for each CU.

###### Late century

Figure 5.5: Overall exposure to climate changes across the lifecycle (x-axis) for 60 salmon CUs in the Fraser River basin (y-axis). The mid-century period (2040-2069) under the RCP 4.5 emissions scenario is included in the main text; **other time periods and emissions scenarios can be viewed here under the tabs.** Points and lines are the median and range in exposure among six different GCMs. CUs are ordered from highest (top) to lowest (bottom) overall exposure. Exposure was calculated as the proportion of the lifecycle that was above or below the thresholds for different climate variables, averaged among variables and life stages for each CU.

##### High Emissions (RCP 8.5)

###### Historical

Figure 5.6: Overall exposure to climate changes across the lifecycle (x-axis) for 60 salmon CUs in the Fraser River basin (y-axis). The mid-century period (2040-2069) under the RCP 4.5 emissions scenario is included in the main text; **other time periods and emissions scenarios can be viewed here under the tabs.** Points and lines are the median and range in exposure among six different GCMs. CUs are ordered from highest (top) to lowest (bottom) overall exposure. Exposure was calculated as the proportion of the lifecycle that was above or below the thresholds for different climate variables, averaged among variables and life stages for each CU.

###### Early century

Figure 5.7: Overall exposure to climate changes across the lifecycle (x-axis) for 60 salmon CUs in the Fraser River basin (y-axis). The mid-century period (2040-2069) under the RCP 4.5 emissions scenario is included in the main text; **other time periods and emissions scenarios can be viewed here under the tabs.** Points and lines are the median and range in exposure among six different GCMs. CUs are ordered from highest (top) to lowest (bottom) overall exposure. Exposure was calculated as the proportion of the lifecycle that was above or below the thresholds for different climate variables, averaged among variables and life stages for each CU.

###### Mid century

Figure 5.8: Overall exposure to climate changes across the lifecycle (x-axis) for 60 salmon CUs in the Fraser River basin (y-axis). The mid-century period (2040-2069) under the RCP 4.5 emissions scenario is included in the main text; **other time periods and emissions scenarios can be viewed here under the tabs.** Points and lines are the median and range in exposure among six different GCMs. CUs are ordered from highest (top) to lowest (bottom) overall exposure. Exposure was calculated as the proportion of the lifecycle that was above or below the thresholds for different climate variables, averaged among variables and life stages for each CU.

###### Late century

Figure 5.9: Overall exposure to climate changes across the lifecycle (x-axis) for 60 salmon CUs in the Fraser River basin (y-axis). The mid-century period (2040-2069) under the RCP 4.5 emissions scenario is included in the main text; **other time periods and emissions scenarios can be viewed here under the tabs.** Points and lines are the median and range in exposure among six different GCMs. CUs are ordered from highest (top) to lowest (bottom) overall exposure. Exposure was calculated as the proportion of the lifecycle that was above or below the thresholds for different climate variables, averaged among variables and life stages for each CU.

#### 5.4 Exposure by life stage and variable

The following figures are expanded version of Figure 4 in the main text, showing the proportion of time above or below the relevant threshold for each stage and climate variable (x -axis) and each CU (y-axis). Here, we also include exposure to sea surface salinity for completeness, but note that this variable was excluded when calculating overall exposure.

##### Low emissions (RCP 4.5)

###### Historical

Figure 5.10: The proportion of time above or below the relevant threshold for each stage and climate variable (x -axis) and each CU (y-axis). The mid-century period (2040-2069) under the RCP 4.5 emissions scenario is included in the main text; **other time periods and emissions scenarios can be viewed here under the tabs.** Stream temperature (temp) and low flow (flow) were assessed for freshwater stages; sea surface temperature (SST) and sea surface salinity (sal) were assessed for marine stages. CUs are ordered from highest (top) to lowest (bottom) exposure to climate changes across the lifecycle and color coded according to species (see overall exposure figure above).

###### Early century

Figure 5.11: The proportion of time above or below the relevant threshold for each stage and climate variable (x -axis) and each CU (y-axis). The mid-century period (2040-2069) under the RCP 4.5 emissions scenario is included in the main text; **other time periods and emissions scenarios can be viewed here under the tabs.** Stream temperature (temp) and low flow (flow) were assessed for freshwater stages; sea surface temperature (SST) and sea surface salinity (sal) were assessed for marine stages. CUs are ordered from highest (top) to lowest (bottom) exposure to climate changes across the lifecycle and color coded according to species (see overall exposure figure above).

###### Mid century

Figure 5.12: The proportion of time above or below the relevant threshold for each stage and climate variable (x -axis) and each CU (y-axis). The mid-century period (2040-2069) under the RCP 4.5 emissions scenario is included in the main text; **other time periods and emissions scenarios can be viewed here under the tabs.** Stream temperature (temp) and low flow (flow) were assessed for freshwater stages; sea surface temperature (SST) and sea surface salinity (sal) were assessed for marine stages. CUs are ordered from highest (top) to lowest (bottom) exposure to climate changes across the lifecycle and color coded according to species (see overall exposure figure above).

###### Late century

Figure 5.13: The proportion of time above or below the relevant threshold for each stage and climate variable (x -axis) and each CU (y-axis). The mid-century period (2040-2069) under the RCP 4.5 emissions scenario is included in the main text; **other time periods and emissions scenarios can be viewed here under the tabs.** Stream temperature (temp) and low flow (flow) were assessed for freshwater stages; sea surface temperature (SST) and sea surface salinity (sal) were assessed for marine stages. CUs are ordered from highest (top) to lowest (bottom) exposure to climate changes across the lifecycle and color coded according to species (see overall exposure figure above).

##### High Emissions (RCP 8.5)

###### Historical

Figure 5.14: The proportion of time above or below the relevant threshold for each stage and climate variable (x -axis) and each CU (y-axis). The mid-century period (2040-2069) under the RCP 4.5 emissions scenario is included in the main text; **other time periods and emissions scenarios can be viewed here under the tabs.** Stream temperature (temp) and low flow (flow) were assessed for freshwater stages; sea surface temperature (SST) and sea surface salinity (sal) were assessed for marine stages. CUs are ordered from highest (top) to lowest (bottom) exposure to climate changes across the lifecycle and color coded according to species (see overall exposure figure above).

###### Early century

Figure 5.15: The proportion of time above or below the relevant threshold for each stage and climate variable (x -axis) and each CU (y-axis). The mid-century period (2040-2069) under the RCP 4.5 emissions scenario is included in the main text; **other time periods and emissions scenarios can be viewed here under the tabs.** Stream temperature (temp) and low flow (flow) were assessed for freshwater stages; sea surface temperature (SST) and sea surface salinity (sal) were assessed for marine stages. CUs are ordered from highest (top) to lowest (bottom) exposure to climate changes across the lifecycle and color coded according to species (see overall exposure figure above).

###### Mid century

Figure 5.16: The proportion of time above or below the relevant threshold for each stage and climate variable (x -axis) and each CU (y-axis). The mid-century period (2040-2069) under the RCP 4.5 emissions scenario is included in the main text; **other time periods and emissions scenarios can be viewed here under the tabs.** Stream temperature (temp) and low flow (flow) were assessed for freshwater stages; sea surface temperature (SST) and sea surface salinity (sal) were assessed for marine stages. CUs are ordered from highest (top) to lowest (bottom) exposure to climate changes across the lifecycle and color coded according to species (see overall exposure figure above).

###### Late century

Figure 5.17: The proportion of time above or below the relevant threshold for each stage and climate variable (x -axis) and each CU (y-axis). The mid-century period (2040-2069) under the RCP 4.5 emissions scenario is included in the main text; **other time periods and emissions scenarios can be viewed here under the tabs.** Stream temperature (temp) and low flow (flow) were assessed for freshwater stages; sea surface temperature (SST) and sea surface salinity (sal) were assessed for marine stages. CUs are ordered from highest (top) to lowest (bottom) exposure to climate changes across the lifecycle and color coded according to species (see overall exposure figure above).

#### 5.5 Exposure across time periods

##### Low emissions (RCP 4.5)

Figure 5.18: Changes in exposure (y-axis), calculated as the proportion of time above or below threshold, for each CU (lines) organized by species (colours), across periods (x-axis; H = historical, E = early century, M = mid century, L = late century). Each panel shows exposure for a specific life stage and climate variable. Each line represents the median exposure among GCMS for a single CU, and the shaded region behind lines indicates the range of exposure outcomes among GCMs for that CU. Exposure is similar for marine stages because all CUs had the same spatial distribution and the monthly resolution of climate projections, combined with less overall variability through time for ocean variables, did not differentiate exposure due to CU-specific life-cycle timing.

##### High emissions (RCP 8.5)

Figure 5.19: Changes in exposure (y-axis), calculated as the proportion of time above or below threshold, for each CU (lines) organized by species (colours), across periods (x-axis; H = historical, E = early century, M = mid century, L = late century). Each panel shows exposure for a specific life stage and climate variable. Each line represents the median exposure among GCMS for a single CU, and the shaded region behind lines indicates the range of exposure outcomes among GCMs for that CU. Exposure is similar for marine stages because all CUs had the same spatial distribution and the monthly resolution of climate projections, combined with less overall variability through time for ocean variables, did not differentiate exposure due to CU-specific life-cycle timing.
